# FcRγ^−^ NK cell induction by specific CMV and expansion by subclinical viral infections in rhesus macaques

**DOI:** 10.1101/2022.05.26.493509

**Authors:** Jaewon Lee, W. L. William Chang, Jeannine M. Scott, Suyeon Hong, Taehyung Lee, Jesse D. Deere, Peter Park, Ellen E. Sparger, Satya Dandekar, Dennis J. Hartigan-O’Connor, Peter A. Barry, Sungjin Kim

## Abstract

Long-lived ‘memory-like’ NK cells, characterized by FcRγ-deficiency and enhanced responsiveness to antibody-bound virus-infected cells, have been found in certain human cytomegalovirus (HCMV)-seropositive individuals. Because humans are exposed to numerous microbes and environmental agents, specific relationships between HCMV and FcRγ-deficient NK cells (also known as g-NK cells) have been challenging to define. Here, we show that a subgroup of rhesus cytomegalovirus (RhCMV)-seropositive macaques possesses FcRγ-deficient NK cells that stably persist and display phenotype resembling human FcRγ-deficient NK cells. Moreover, these macaque NK cells resembled human FcRγ-deficient NK cells with respect to functional characteristics, including enhanced responsiveness to RhCMV-infected target in an antibody-dependent manner and hypo-responsiveness to tumor and cytokine stimulation. These cells were not detected in specific-pathogen-free (SPF) macaques free of RhCMV and six other viruses; however, experimental infection of SPF animals with RhCMV strain UCD59, but not RhCMV strain 68-1 or SIV, led to induction of FcRγ-deficient NK cells. In non-SPF macaques, co-infection by RhCMV with other common viruses was associated with higher frequencies of FcRγ-deficient NK cells. These results support a causal role for specific cytomegalovirus strain(s) in the induction of FcRγ-deficient NK cells, and suggest that co-infection by other viruses further expands this memory-like NK cell pool.

## Introduction

Although NK cells have been traditionally considered part of the innate immune system, several recent studies have revealed subsets of NK cells with adaptive immune features, including antigen-specific memory responses (1-8). However, since NK cells express only germline-encoded receptors, they presumably cannot mount antigen-specific memory responses to diverse targets through direct recognition.

We discovered a distinct subset of human NK cells that displays adaptive immune features, including clonal-like expansion and long-term persistence (9-11). This novel NK cell subset, named g-NK cells, is characterized by deficiency in FcRγ, a signaling adaptor normally associated with the IgG Fc receptor CD16, either as a homodimer or as a heterodimer with another adaptor CD3ζ (12). The deficiency appears to result from DNA hypermethylation in the FcRγ gene, along with other epigenetic modifications (13), which are associated with altered expression of multiple proteins, including SYK tyrosine kinase. Importantly, human g-NK cells exhibit enhanced abilities to produce cytokines and expand in number upon interaction with cells infected with viruses (e.g., HCMV and influenza virus) in the presence of virus-specific antibodies via the action of CD16 (11). Additionally, we have determined g-NK cells can constitute as much as 85% of circulating NK cells, thereby surpassing the number of circulating memory CD8^+^ T cells in certain individuals (9, 10). Therefore, g-NK cells have heightened potential for protective immune responses to a broad spectrum of viral infections through antibody-dependent memory-like effector functions (11, 14-17).

Despite their important adaptive immune features, little is known about how this memory-like NK cell pool is induced and shaped within individuals. To date, g-NK cells have been detected at a wide range of frequencies in approximately one-third of healthy people (9, 10, 14, 16, 18, 19). Epidemiological analyses indicate that the presence of these cells is associated with seropositivity for HCMV (10, 14, 18). However, because humans are naturally exposed to numerous microbes and environmental agents, including other viruses, and because g-NK cells were detectable in only a subgroup of HCMV-seropositive individuals, it has been challenging to determine the specific role HCMV might play in the induction and/or expansion of g-NK cells. Interestingly, HCMV-seropositivity has also been associated with ‘adaptive’ NKG2C^+^ NK cells in a manner similar to g-NK cells (20-25). Establishing whether there is a causal relationship between HCMV infection and g-NK cell development will be an important step toward delineating the specific factors and processes by which this memory-like NK cell pool is induced and shaped through expansion. However, a recent attempt to induce FcRγ-deficient NK cells in mice by experimental infection with murine CMV was unsuccessful (11), leaving no current mouse model available for the study of g-NK cells.

In this study, we found that a subgroup of non-SPF rhesus macaques with naturally acquired RhCMV infection possessed FcRγ-deficient NK cells (hereafter called FcRγ^−^ NK cells) that displayed phenotypic and functional characteristics resembling human g-NK cells. Importantly, these cells were not detected in SPF animals. However, experimental RhCMV infection of SPF animals led to the induction of FcRγ^−^ NK cells in a strain-specific manner. Serological analysis of non-SPF animals revealed that subclinical infections by other common viruses can contribute to the expansion of this adaptive, memory-like NK cell pool.

## Results

### Identification and long-term persistence of FcRγ^−^ NK cells in rhesus macaques

To explore a non-human primate model for the study of g-NK cells, we examined the expression of CD16-associated FcRγ and CD3ζ adaptors in NK cells from peripheral blood samples of rhesus macaques. Since previous studies have shown that the majority of macaque NK cells express NKG2A (5, 26, 27), we initially used this marker to identify NK cells as CD3ε^−^CD14^−^CD20^−/dim^ and NKG2A^+^. Using this gating strategy, we observed that all CD3ε^−^CD14^−^CD20^−/dim^NKG2A^+^ cells expressed CD3ζ in all macaque subjects examined, but several macaques (e.g. RM^#^663) harbored a distinct subset that was deficient in FcRγ (Fig. 1A). Throughout this investigation, we used anti-FcRγ mAb (1D6), which allowed for much sharper resolution between these macaque FcRγ^−^ NK cells and FcRγ-expressing conventional NK cells compared to the polyclonal antibodies used in previous studies (9-11, 28, 29).

**Figure 1.**
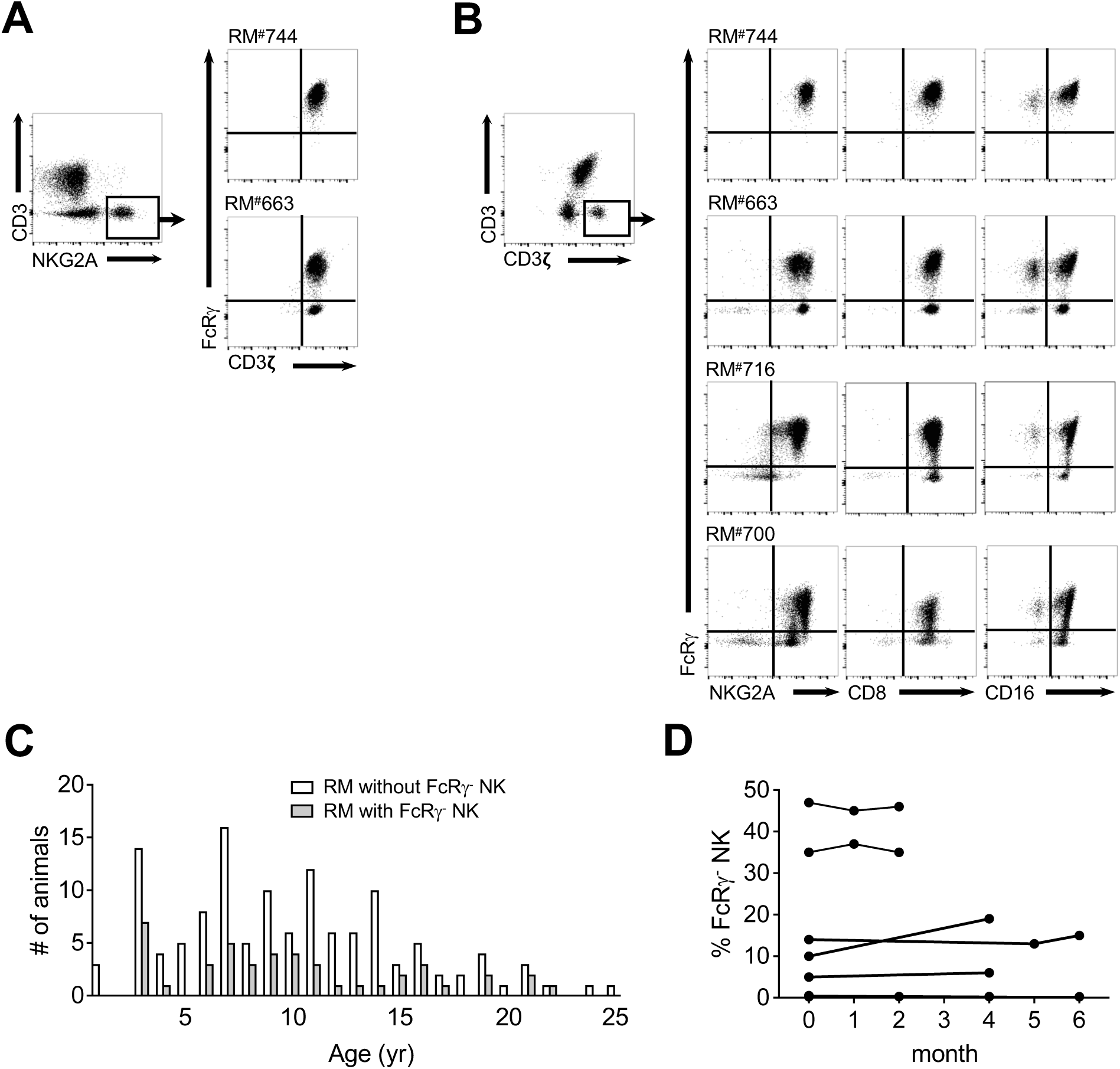
FcRγ^−^ NK cells are present in a subgroup of rhesus macaques and stably persist. Macaque PBMCs were analyzed by flow cytometry for the presence of FcRγ^−^ NK cells. Live lymphocytes were gated as CD3ε^−^CD14^−^CD20^−/dim^NKG2A^+^(A) or CD3ε^−^CD14^−^CD20^−/dim^CD3ζ^+^(B). Dot plots show presence or absence of FcRγ^−^ NK cells along with expression of indicated markers from several representative macaques. (C) PBMCs from individual macaques (*n*=128) were analyzed to determine the presence of FcRγ^−^ NK cells among the CD3ε^−^CD14^−^CD20^−/dim^CD3ζ^+^CD16^+^ NK cell population for each sample. Bar graph shows the number of surveyed animals with (grey bar) and without (white bar) FcRγ^−^ NK cells at specified ages. (D) Longitudinal monitoring of FcRγ^−^ NK cell subset. Shown are frequencies of FcRγ^−^ NK cell subsets among the CD3ε^−^CD14^−^CD20^−/dim^CD3ζ^+^CD16^+^ NK cell population in PBMC samples collected for up to 6 months from 7 individual macaques.

Because many g-NK cells in humans do not express NKG2A (10, 17), gating strategies that utilize NKG2A might not include the entire FcRγ^−^ NK cell population in macaques. Therefore, we considered alternative markers to identify all NK cells in rhesus macaques. Since all human NK cells express CD3ζ (9), and because this adaptor appeared to be expressed exclusively by NK cells and T cells in both humans and macaques (Fig. 1A and data not shown), we examined FcRγ expression in the CD3ε^−^CD14^−^CD20^−/dim^CD3ζ^+^ cell population to determine the frequency of FcRγ^−^ NK cells among total circulating NK cells in macaques. Indeed, this gating strategy revealed that some animals (e.g. RM^#^716 and ^#^700) had readily detectable, albeit often at low frequency, FcRγ^−^ NK cells that expressed little or low levels of NKG2A in addition to FcRγ^−^ NK cells that expressed high levels of NKG2A (Fig. 1B). The majority of these

CD3ε^−^CD14^−^CD20^−/dim^CD3ζ^+^NKG2A^−/dim^ FcRγ^−^ cells expressed NK cell-associated markers CD8 and CD16 (30), and the effector molecule granzyme B (Fig. 1B and Supplemental Fig. 1), indicating that NKG2A^−/dim^ NK cells that are also FcRγ-deficient exist in certain macaques. Although the CD3ε^−^CD14^−^CD20^−/dim^CD3ζ^+^ gating strategy was more inclusive of FcRγ^−^ NK cells, the gated population appeared to contain a small population of cells that did not express NK cell-associated proteins, including CD16 (Supplemental Fig. 1). Therefore, we modified the gating strategy to include CD16 expression to more accurately assess the frequency of FcRγ^−^ NK cells with a final profile defined as CD3ε^−^CD14^−^CD20^−/dim^CD3ζ^+^CD16^+^ FcRγ^−^ NK cells. The resulting analysis of a large cohort of non-SPF macaques (*n*=128) using this gating strategy showed that approximately one-third of the macaques had a distinct population of FcRγ^−^ NK cells (Fig. 1C). The incidence and frequencies of FcRγ^−^ NK cells did not show notable association with age or the gender of the animal subjects (Fig. 1C and data not shown). These data demonstrate that a subgroup of rhesus macaques possesses FcRγ^−^ NK cells, similar to the observed incidence of g-NK cells in humans (9, 14, 18).

Longitudinal studies showed that the frequencies of circulating FcRγ^−^ NK cells were nearly constant over a period of 4 to 6 months (Fig. 1D). In contrast, macaques that initially had no detectable FcRγ^−^ NK cells showed no appearance of such cells during this period. These data indicate that the size of FcRγ^−^ NK cell pool is maintained at a relatively stable level.

### Phenotypic characteristics of FcRγ^−^ NK cells

It has been shown that several cell surface markers and intracellular proteins, such as NKp46 and SYK, are differentially expressed between human g-NK and conventional NK cells (9-11, 14, 17). Consistent with this phenotypic difference in human NK cells, macaque FcRγ^−^ NK cells showed lower levels of both NKp30 and NKp46 compared to conventional NK cells (Fig. 2A). Also resembling human NK cell populations (11), the expression of adhesion molecules PECAM and CD49F was lower, while CD2 was higher in FcRγ^−^ NK cells compared to conventional NK cells (Fig. 2A).

**Figure 2.**
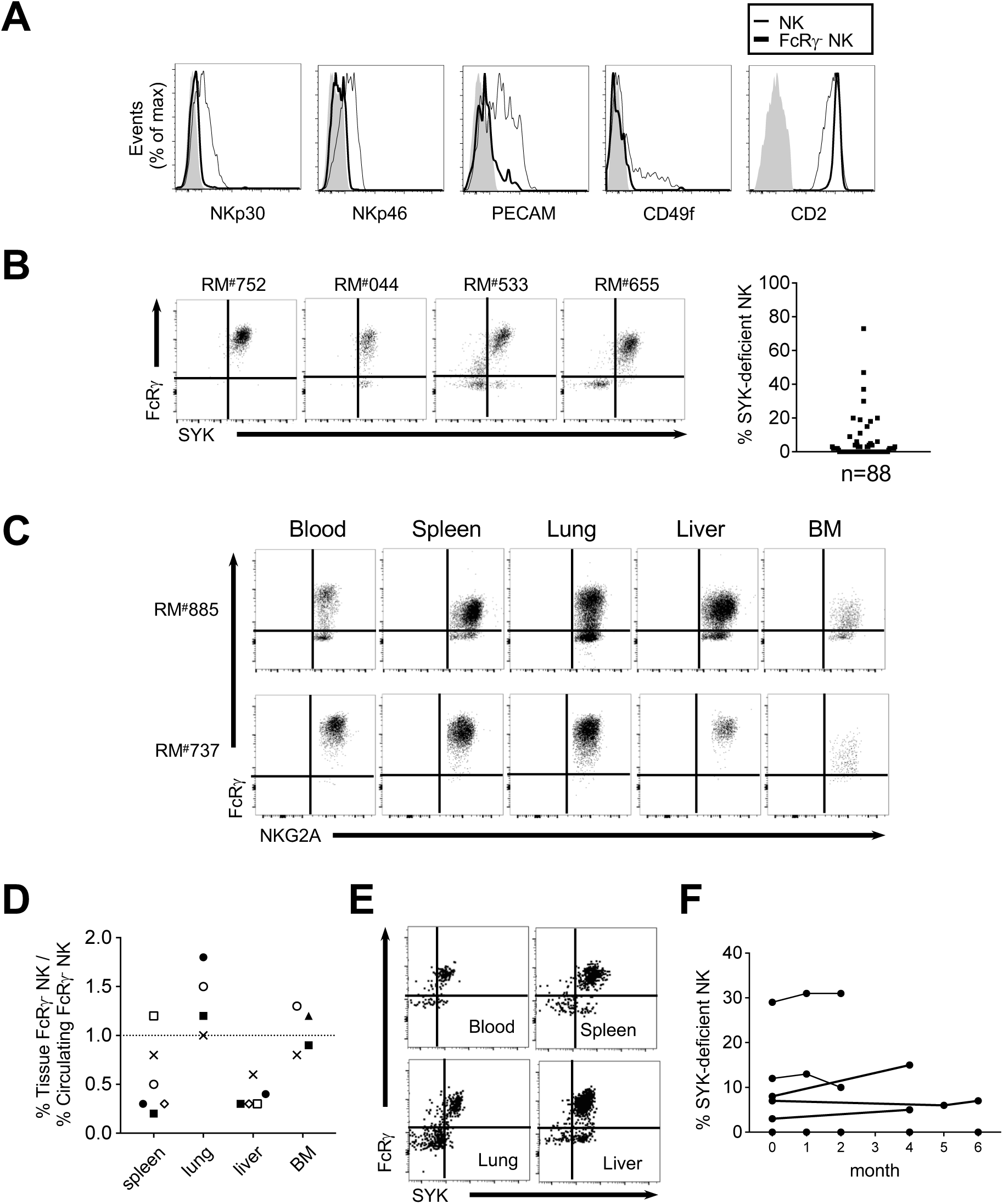
Phenotypic properties, tissue distribution and long-term persistence of SYK-deficient NK cells. (A) Histograms show expression of indicated surface markers on FcRγ^−^ NK cells (bold line), conventional NK cells (thin line), and staining controls (grey shaded) in the CD3ε^−^ CD14^−^CD20^−/dim^CD3ζ^+^CD16^+^ NK cell population. Data are representative of seven macaque PBMC samples. (B) Expression of FcRγ and SYK in the CD3ε^−^CD14^−^CD20^−/dim^CD3ζ^+^CD16^+^ NK cell population from PBMCs of four representative macaques. Dot graph shows the frequencies of SYK-deficient cells among the NK cell population in PBMCs, with each data point representing an individual animal (*n*=88). (C) FcRγ and NKG2A expression of NK cell population in PBMCs, spleen, lung, liver, and bone marrow (BM) from two representative macaques. (D) Comparative ratio of frequencies of tissue-resident FcRγ^−^ NK cells to circulating FcRγ^−^ NK cells in each animal. Each symbol represents data points from the same macaque. (E) FcRγ and SYK expression in NK cells from peripheral blood, spleen, lung, and liver samples from a representative macaque. (F) Longitudinal monitoring of the SYK-deficient NK cell subset. Shown are frequencies of SYK-deficient NK cells among the CD3ε^−^CD14^−^CD20^−/dim^CD3ζ^+^CD16^+^ NK cell population in PBMC samples collected for up to 6 months from 7 individual macaques.

Moreover, while macaque conventional NK cells expressed SYK at uniform levels, FcRγ^−^ NK cells showed variable levels of SYK among different animals; i.e., FcRγ^−^ NK cells displayed near complete or partial SYK-deficiency in some animals, while displaying near normal SYK expression in other animals (Fig. 2B). Overall, FcRγ^−^ NK cells showed reduced, yet heterogeneous, expression of SYK, and SYK-deficient NK cells were found in approximately one quarter of the animals examined. Taken together, these data show that macaque FcRγ^−^ NK cells and human g-NK cells have several phenotypic characteristics in common.

Analysis of tissue specimens showed that these cells were present in spleen, lungs, liver, and bone marrow in macaques that had FcRγ^−^ NK cells in peripheral blood (Fig. 2C). In contrast, macaques that lacked FcRγ^−^ NK cells in peripheral blood consistently lacked detectable FcRγ^−^ NK cells in these tissues. Interestingly, compared to peripheral blood, the relative frequency of FcRγ^−^ NK cells was higher in lung, lower in spleen and liver, and comparable in bone marrow (Fig. 2D). SYK-deficient NK cells were also present in these tissues from macaques with SYK-deficient NK cells in peripheral blood (Fig. 2E). These data demonstrated that FcRγ^−^ NK cells, as well as SYK-deficient NK cells, were widely distributed in the body at variable frequencies depending on the tissues. Longitudinal studies showed that the frequencies of SYK-deficient NK cells were also nearly constant during the follow-up period (Fig. 2F), again resembling the persistent presence of SYK-deficient NK cells observed in humans. (11).

### Enhanced antibody-dependent activation of FcRγ^−^ NK cells against virus-infected cells

One of the defining features of human g-NK cells is their enhanced ability to respond to virus-infected target cells in the presence of virus-specific antibodies (10, 11, 14-17). To determine whether macaque FcRγ^−^ NK cells also exhibit this functional property, we examined NK cell intracellular cytokine expression by flow cytometry after stimulation with rhesus fibroblasts (Telo-RF) that were either RhCMV- or mock-infected in the presence or absence of RhCMV-seropositive plasma. In response to mock-infected target cells, both FcRγ^−^ and conventional NK cells revealed little or no production of TNF-α regardless of the presence of seropositive plasma (Fig. 3A).

**Figure 3.**
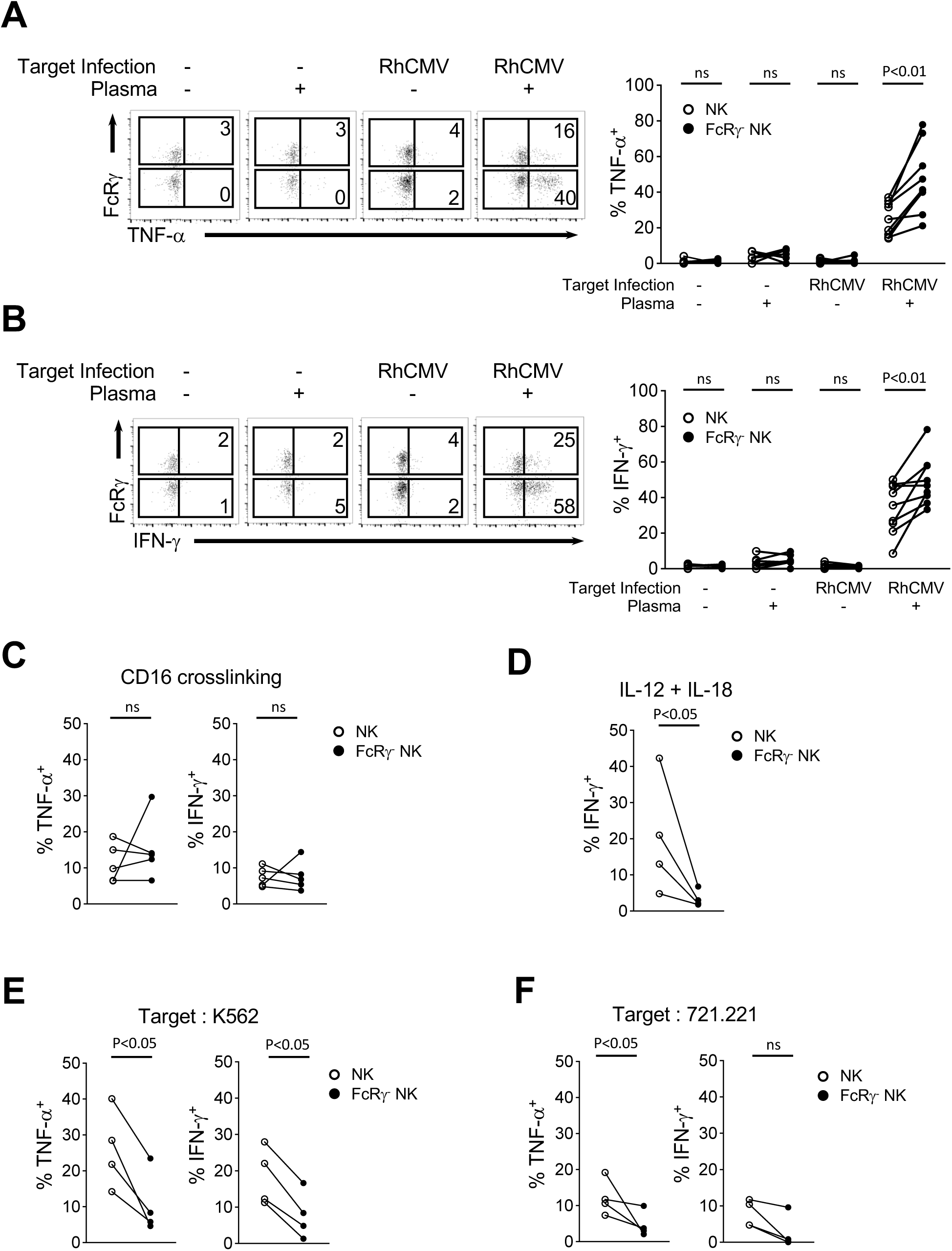
FcRγ^−^ NK cells show enhanced effector function against virus-infected target cells in the presence of virus-specific antibodies. (A and B) Macaque PBMCs were co-cultured with mock- or RhCMV-infected cells in the presence or absence of RhCMV-specific plasma as indicated. Induction of TNF-α (A) and IFN-γ (B) by conventional NK and FcRγ^−^ NK cells were assessed by flow cytometry. Numbers represent the relative percentage of TNF-α^+^ or IFN-γ^+^ NK cells among each NK cell subset. Line graphs on the right show the percentages of responding conventional NK (○) or FcRγ^−^ NK (●) cells. Circles connected by a line designate the same macaque sample. (C-F) Percentages of conventional NK (○) or FcRγ^−^ NK (●) cells that expressed TNF-α or IFN-γ following stimulation with CD16 crosslinking (C), IL-12 and IL-18 (D), K562 (E) or 721.221 (F) tumor target cells. Statistical analyses were performed using nonparametric Mann-Whitney tests. ns, not significant.

Similarly, neither FcRγ^−^ nor conventional NK cells produced appreciable amounts of TNF-α in response to RhCMV-infected cells in the absence of RhCMV-seropositive plasma. However, the addition of seropositive plasma led to a dramatic increase in frequency of TNF-α expressing FcRγ^−^ NK cells at levels significantly higher than that of TNF-α expressing conventional NK cells (p<0.01) (Fig. 3A). In contrast, the addition of RhCMV seronegative plasma did not elicit these effects (data not shown). Additionally, significantly higher frequencies of IFN-γ expressing cells were observed within the FcRγ^−^ NK cell subset compared to conventional NK cells when both RhCMV-infected cells and seropositive plasma were present (Fig. 3B). These results showed that, compared to conventional NK cells, FcRγ^−^ NK cells exhibited enhanced responses to RhCMV-infected cells in an antibody-dependent manner. However, macaque FcRγ^−^ NK cells did not show higher cytokine production compared to conventional NK cells when stimulated with immobilized anti-human CD16 mAb (3G8) (Fig. 3C).

It has been shown that human g-NK cells respond poorly to IL-12 and IL-18 co-stimulation (14), consistent with the observed lower levels of transcripts encoding IL-12Rb2 and IL-18RAP receptor subunits determined via transcriptome analyses (11, 14). Similarly, treatment of macaque FcRγ^−^ NK cells with these cytokines resulted in much lower frequencies of IFN-γ-expressing cells compared to conventional NK cells (p<0.05) (Fig. 3D). Finally, we have previously shown that human g-NK cells respond poorly to certain tumor cells in the absence of tumor cell-specific antibodies (9). To compare the functional responsiveness of macaque FcRγ^−^ NK cells and conventional NK cells to tumor targets, we examined cytokine expression following stimulation with human K562 or 721.221 tumor cells. Exposure of FcRγ^−^ NK cells to either target cell line resulted in lower frequencies of TNF-α and IFN-γ-expressing cells compared to conventional NK cells (p<0.05) (Fig. 3E and 3F). These data indicate that FcRγ^−^ NK cells are hypo-responsive to stimuli other than antibody-dependent stimulation.

### Induction of FcRγ^−^ NK cells following experimental infection with a specific strain of RhCMV

Considering the association between HCMV serostatus and g-NK cells (10, 11, 14), we hypothesized that RhCMV infection may play a role in the induction of FcRγ^−^ NK cells in macaques. To explore this possibility, we first determined whether FcRγ^−^ NK cells are present in SPF macaques that were raised in an outdoor environment free from RhCMV as well as six other viruses that include simian foamy virus (SFV), herpes B virus (BV) rhesus monkey rhadinovirus (RRV), type D simian retrovirus (SRV), simian immunodeficiency virus (SIV) and simian T-lymphotropic virus (STLV). These SPF animals were confirmed seronegative for all seven viruses, and ranged in age from 2 to 13 years, with the median age of 5 years. Importantly, analysis of peripheral blood samples from these SPF macaques showed that none had detectable FcRγ^−^ NK cells, and all NK cells maintained uniformly high levels of NKG2A and SYK (Fig. 4A and 4B). Thus, these data suggest that regardless of age, the presence of FcRγ^−^ NK cells in rhesus macaques requires infection with at least one of these seven viruses.

**Figure 4.**
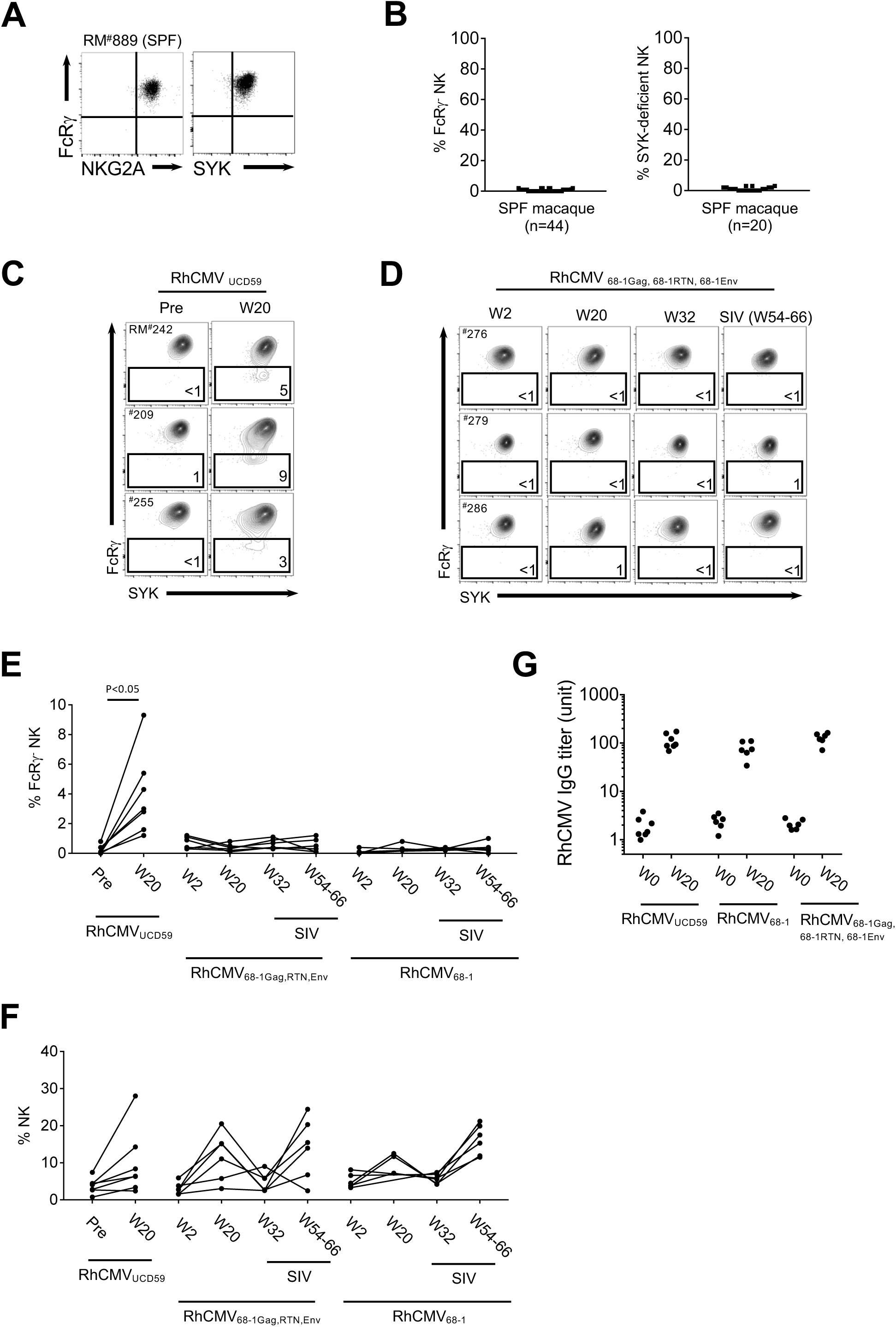
FcRγ^−^ NK cells are induced in SPF macaques following primary infection with a specific RhCMV strain. (A) Expression of NKG2A and SYK with respect to FcRγ in the CD3ε^−^CD14^−^CD20^−/dim^CD3ζ^+^CD16^+^ NK cell population from a representative SPF macaque. (B) Frequencies of FcRγ^−^ and SYK-deficient NK cells in PBMCs of SPF macaques. (C) FcRγ and SYK expression in NK cells from PBMC samples from three SPF macaques before (Pre) and 20 weeks (W20) after RhCMV_UCD59_ infection. (D) FcRγ and SYK expression in NK cells from PBMC samples from three SPF macaques at 2 weeks (W2), 20 weeks (W20) and 32 weeks (W32) after infection with combination of RhCMV_68-1Gag_, RhCMV_68-1RTN_, and RhCMV_68-1Env_. These macaques were subsequently challenged with SIV at 36 weeks after initial RhCMV infection. Dot plots show flow cytometric analysis of FcRγ and SYK expression in NK cells obtained before (W32) and between 22-34 weeks post-SIV infection. (E) Frequencies of FcRγ^−^ deficient NK cells are shown before and after infection by indicated strains of RhCMV and SIV from all macaques. (F) NK cell frequencies are shown before and after infection by indicated strains of RhCMV and SIV. (G) Plasma IgG titers for RhCMV from macaques before (W0) and 20 weeks after infection by indicated strains of RhCMV.

We next sought to examine the role of RhCMV infection in the induction of FcRγ^−^ NK cells through retrospective analysis of PBMC samples that had been collected from SPF animals (*n*=7) pre- and post-infection with RhCMV_UCD59_, a strain originally isolated from a non-SPF macaque (31). While the analysis of baseline samples of uninfected SPF animals showed no detectable FcRγ^−^ NK cells initially, at 20 weeks post-infection, a noticeable subset of NK cells that was deficient for FcRγ was observed in several animals, albeit at low frequencies (Fig. 4C and 4E), providing evidence for a role of RhCMV infection in the induction of FcRγ^−^ NK cells. We also conducted retrospective analysis of PBMCs collected from a group of SPF animals (*n*=6) that had been experimentally infected with a mixture of three variants of RhCMV_68-1_, a prototypic fibroblast-tropic strain of RhCMV that has been widely used for experimental infection of macaques, including vaccine studies (32-34). Intriguingly, FcRγ^−^ NK cells were not detected in any of the PBMC samples collected at 2, 20, or 32 weeks post-infection with these RhCMV_68-1_ variants (Fig. 4D and 4E). In addition, we analyzed PBMCs collected from a separate group of SPF macaques (*n*=6) that were infected with RhCMV_68-1_ alone. Again, no animals infected with RhCMV_68-1_ revealed detectable FcRγ^−^ NK cells for up to 32 weeks post-infection (Fig. 4E). As part of a vaccine study, these animals were challenged with SIV starting from 36 weeks after infection with RhCMV_68-1_ or RhCMV_68-1_ variants, and the subsequent analysis of PBMCs collected between 22 - 34 weeks after SIV infection showed no evidence of FcRγ^−^ NK cells (Fig. 4D and 4E). Taken together, these results indicate that unlike RhCMV_UCD59_, infection with RhCMV_68-1_, RhCMV_68-1_ variants or SIV did not induce FcRγ^−^ NK cells during the indicated time periods.

Importantly, the RhCMV strain-specific effect we observed was not due to a difference in the ability of these viruses to induce a global expansion of NK cells. Indeed, infection with RhCMV_68-1_, RhCMV_68-1_ variants, and SIV led to the expansion of NK cells at levels comparable to those observed following infection with RhCMV_UCD59_ (Fig. 4F). Moreover, infection with RhCMV_68-1_ and RhCMV_68-1_ variants elicited RhCMV-specific IgG at concentrations comparable to those observed following infection with RhCMV_UCD59_ (Fig. 4G), likely reflecting productive long-term infection by these strains.

### Association of FcRγ^−^ NK cells with natural infection by RhCMV and other viruses

Considering the strain-specific effect observed with experimental RhCMV infection, we examined the potential relationship between natural RhCMV infection and FcRγ^−^ NK cells in non-SPF macaques. Serological analysis of plasma samples for RhCMV-specific antibodies showed that the majority of non-SPF macaques were seropositive for RhCMV (Fig. 5A). Among the RhCMV-seropositive macaques examined, approximately one-third had detectable FcRγ^−^ NK cells in their peripheral blood, while this subset was detected in only one RhCMV-seronegative macaque. Both the incidence and frequency of FcRγ^−^ NK cells were significantly higher in the RhCMV-seropositive group than in the RhCMV-seronegative group (p<0.01) (Fig. 5A), indicating the association of FcRγ^−^ NK cells with naturally acquired RhCMV infection.

**Figure 5.**
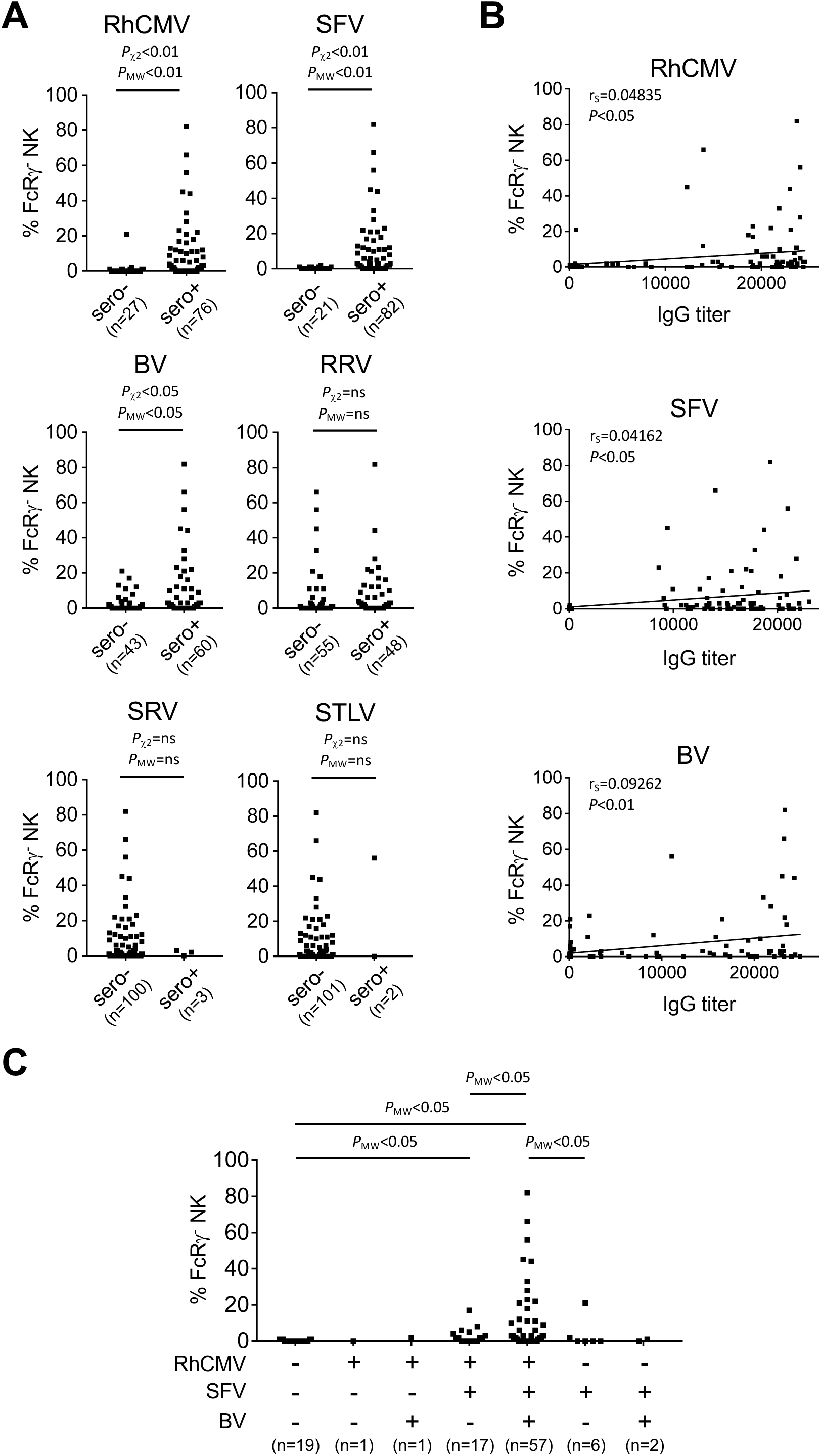
Association of FcRγ^−^ NK cells with infection by RhCMV, SFV and BV. (A) Frequencies of FcRγ^−^ NK cells within individual non-SPF macaques are grouped according to IgG serostatus for infection of RhCMV, SFV, BV, RRV, SRV and STLV. (B) FcRγ^−^ NK cell frequencies were plotted according to plasma IgG titers for each animal for indicated viruses. (C) Frequencies of FcRγ^−^ NK cells within individual macaques grouped according to their IgG serological statuses with respect to 3 common viruses (RhCMV, SFV and BV). Statistical analyses for the incidence and frequencies of FcRγ^−^ NK cells were performed using chi-square tests (P_χ2_) and nonparametric Mann-Whitney tests (P_MW_), respectively. Correlation analyses were performed using Spearman’s rank test. ns, not significant.

Since the frequencies of FcRγ^−^ NK cells in the SPF macaques experimentally infected with RhCMV_UCD59_ were generally lower than those detected in non-SPF RhCMV-seropositive macaques (Fig. 4C and Fig. 5A), and since non-SPF animals are exposed to potentially many other viruses, microbes, or environmental agents, we hypothesized that these additional exposures might contribute to the higher frequencies of FcRγ^−^ NK cells in these animals. To investigate this possibility, we assessed the levels of plasma antibodies specific for each of the six other viruses known to be absent in SPF macaques. Serologic results showed that, similar to RhCMV, the majority of macaques were seropositive for SFV. Approximately one half of the animals were seropositive for BV and RRV, only a small fraction of macaques was seropositive for SRV and/or STLV, and none were seropositive for SIV (Fig. 5A and data not shown). Thus, SFV, BV and RRV were also relatively common in our non-SPF cohorts. Interestingly, FcRγ^−^ NK cells were detected in a subgroup of SFV-seropositive macaques, but not in SFV-seronegative macaques, indicating a positive correlation between prior SFV infection and the presence of FcRγ^−^ NK cells (p<0.01). In addition, the incidence of FcRγ^−^ NK cells was slightly higher in the BV-seropositive group than in the BV-seronegative group (p<0.05), with the seropositive group showing significantly higher frequencies of FcRγ^−^ NK cells (p<0.05). In contrast, neither the incidence nor frequency of FcRγ^−^ NK cells differed between the RRV-seropositive vs. -seronegative animals. Potential associations with SRV or STLV could not be reliably evaluated due to the limited number of animals that were seropositive for these viruses. The frequencies of FcRγ^−^ NK cells showed positive correlation with plasma IgG titers against RhCMV, SFV and BV (Fig. 5B). These data indicate that infection with SFV and/or BV, as well as RhCMV, but not RRV, may be associated with higher incidence and frequency of FcRγ^−^ NK cells in non-SPF macaques.

To explore the possibility that co-infection of RhCMV with SFV and/or BV might influence the incidence and frequency of FcRγ^−^ NK cells, we categorized the animals in our cohorts according to their serological status for each of these three viruses. Notably, all macaques with FcRγ^−^ NK cells, except for one, were seropositive for both RhCMV and SFV, regardless of BV-seropositivity (Fig. 5C and Supplemental Fig. 2). Thus, the data did not allow for further dissection of potential impact of RhCMV and SFV on the incidence and frequency of FcRγ^−^ NK cells. The single exception among the animals with FcRγ^−^ NK cells was an individual that was seropositive for both SFV and RRV (Supplemental Fig. 2); however, none of the animals that were seropositive for either SFV or RRV and seronegative for RhCMV had detectable FcRγ^−^ NK cells (Supplemental Fig. 2). Importantly, although our epidemiologic analysis did not reveal specific association with a particular virus, there was a trend toward higher frequencies of FcRγ^−^ NK cells in the animals that were seropositive for multiple viruses (Fig. 5C). For instance, all animals with relatively high frequencies (>20%) of FcRγ^−^ NK cells were seropositive for BV as well as RhCMV and SFV, indicating that these animals had been exposed to all three viruses. Taken together, these data indicate that co-infection with RhCMV and other common viruses, such as SFV and BV, corelates with higher frequencies of FcRγ^−^ NK cells in non-SPF animals.

## Discussion

Here, we demonstrate that a subgroup of non-SPF macaques with RhCMV sero-positivity possess FcRγ^−^ NK cells that resemble human g-NK cells with adaptive features. Macaque FcRγ^−^ NK cells showed phenotypic characteristics similar to g-NK cells, including reduced expression of natural cytotoxicity receptors, NKp30 and NKp46 (9). In addition, FcRγ^−^ NK cells displayed long-term persistence in macaques. Furthermore, consistent with functional characteristics of g-NK cells, macaque FcRγ^−^ NK cells were hypo-responsive to tumor and cytokine stimulation but were highly responsive to RhCMV-infected cells in an antibody-dependent manner. We speculate that FcRγ^−^ NK cells may play a specialized role in responding to conditions of chronic, reactivated, or recurrent viral infection where pathogen-specific antibodies are available.

Importantly, we observed that SPF animals free of RhCMV and six other viruses did not have detectable FcRγ^−^ NK cells, indicating that one or more viral infection is required for the induction of these cells. The appearance of FcRγ^−^ NK cells in SPF macaques following RhCMV_UCD59_ infection provides strong support for a causal relationship between RhCMV infection and the induction of FcRγ^−^ NK cells, and presumably an analogous causal relationship between HCMV and g-NK cells in humans. The lack of detectable FcRγ^−^ NK cells in SPF macaques following infection with RhCMV_68-1_ and its variants suggests that there is an RhCMV strain-specific effect. The observation that RhCMV_UCD59_ and RhCMV_68-1_ strains were able to elicit comparably high levels of RhCMV-specific IgG suggests that both strains resulted in productive long-term infection. Consistent with this, it has been shown that RhCMV_68-1_ can disseminate and persist in many tissues throughout the body following experimental infection (32, 34, 35). Thus, it is unlikely that the appearance of FcRγ^−^ NK cells following RhCMV_UCD59_ infection is simply due to expansion of pre-existing FcRγ^−^ NK cells that might be present at very low frequencies in these SPF macaques before experimental viral infection. However, considering that one of RhCMV-seronegative macaques had readily detectable FcRγ^−^ NK cells, our data do not exclude the possibility that infection with virus(es) other than RhCMV can also lead to the induction of this NK cell subset. It is also possible that this animal was indeed infected with RhCMV but serological testing did not detect RhCMV-specific antibodies, similar to the reported lack of serological detection of HCMV infection in certain individuals (10, 36). Nonetheless, our retrospective study provides evidence that infection with a specific RhCMV strain is sufficient to induce FcRγ^−^ NK cells, albeit at low frequencies, in the absence of naturally infecting RhCMV and six other viruses in SPF macaques.

The RhCMV strain-specific effect observed in this study provides a potential explanation for why FcRγ^−^ NK cells have been detected in only certain cytomegalovirus-seropositive macaques and humans. Additionally, differences between strains of RhCMV may also account for the apparent lack of association between FcRγ^−^ NK cells and CMV reactivation following hematopoietic stem cell transplantation in rhesus macaques (37). Nucleotide sequence analysis indicates that the main difference between RhCMV_UCD59_ and RhCMV_68-1_ lies in the UL/b’ region that contains several genes involved in regulating host cell tropism and immune evasion (31, 38, 39), suggesting these genes may be important in the induction of FcRγ^−^ NK cells. Prospective infection studies utilizing SPF macaques and RhCMV strains with variations in these genes will be instrumental for delineating viral factors and cellular processes that lead to the induction of FcRγ^−^ NK cells.

Concerning the wide variation in observed frequencies of macaque FcRγ^−^ NK cells, our results from non-SPF macaques suggest that co-infection of RhCMV with other viruses, such as SFV and BV, can lead to further expansion of these cells. In this regard, we have previously demonstrated that g-NK cells preferentially expand over conventional NK cells *in vitro* in response to influenza virus-infected cells, as well as HCMV-infected cells, in a manner dependent on virus-specific antibodies (11), supporting the idea that persistent or recurrent infection by various viruses may drive antibody-dependent expansion of FcRγ^−^ NK cells *in vivo*. Two recent studies have reported that FcRγ^−^ NK cells are present in the majority of macaques from two specific cohorts (28, 29) at what appears to be generally higher frequencies than we observed in our cohort. It is unclear whether the apparent discrepancy is due to the use of different antibodies (monoclonal vs. polyclonal) for FcRγ detection or to a difference in housing environment and/or genetic make-up of macaques. Along this line, the expression of NKp46 was not altered when polyclonal anti-FcRγ was used (29), in contrast to the reduced expression of this activation receptor on FcRγ^−^ NK cells in our study.

Nonetheless, these studies suggested that FcRγ^−^ NK cells can expand in non-SPF macaques following SIV infection, further supporting the role of other viral infections in the expansion of these cells.

Experimental infection of SPF macaques with viruses establishing persistent infection, in combination with RhCMV variants, will be useful for determining the potential contributions of these viruses to the expansion of FcRγ^−^ NK cells. These infection models will be also useful for determining the kinetics of FcRγ^−^ NK cell expansion (19), as well as for evaluating the roles of viral genes (40-46), such as those that are involved in modulating NKG2C-mediated activation of ‘adaptive’ NK cells (47, 48). Based on our observations from the experimental RhCMV infection studies, and the incidence of FcRγ^−^ NK cells in non-SPF macaques, we propose a two-step model (induction and expansion) in which the FcRγ^−^ NK cell population is initially induced by infection with a strain of CMV that bears, as of yet, undefined properties, perhaps leading to epigenetic changes in a small number of NK cells. Subsequent superinfection or reactivation of CMV, and/or recurrent infection by other pathogens where specific antibodies are present could then drive expansion of these FcRγ^−^ NK cells, thereby shaping the antibody-dependent memory-like NK cell pool in an ongoing manner.

## Methods

### Ethics Statement

All studies conducted at the California National Primate Research Center (CNPRC) were approved in advance by the University of California Davis (UC Davis) Institutional Animal Care and Use Committee (approval numbers 16779, and 17880). UC Davis has an Animal Welfare Assurance on file with the National Institutes of Health Office of Laboratory Animal Welfare, and is fully accredited by the Association for the Assessment and Accreditation of Laboratory Animal Care International. All animals had visual and auditory access to other macaques 24 hours per day, and were fed a balanced commercial macaque chow (Purina Mills, Gray Summit, MO) twice daily with fresh produce twice weekly, and free access to water 24 hours per day. Supplemental food was provided when clinically indicated. Environmental enrichment was provided daily, including manipulanda (forage boards, mirrors, puzzle feeders) and novel foodstuffs. When immobilization was necessary, animals were administered 10 mg/kg body weight ketamine-HCl (Parke-Davis, Morris Plains, NJ). The Guidelines for Humane Euthanasia of Animals on Projects (GHEAP) at the CNPRC were followed by the clinical veterinary staff to determine if euthanasia was indicated before the planned endpoint. GHEAP provides guidelines for selecting an endpoint that reduces animal pain and/or distress, while still meeting research objectives. All possible efforts were made to minimize pain and discomfort, including those associated with SIV-associated disease. SIV infection of susceptible macaques can produce a progressive fatal immunodeficiency disease characterized by hematologic abnormalities, lymphocyte depletion, diarrhea, weight loss and cachexia, and infection with opportunistic pathogens. Analgesics were given to minimize pain and discomfort at the discretion of the CNPRC veterinary staff and nutritional supplements were administered as necessary.

### Rhesus Macaques and Viral Infections

SPF cohort of rhesus macaques (*Macaca mulatta*) were maintained as free of RhCMV, SFV, BV, RRV, SIV, SRV and STLV. For RhCMV infection study, a group of animals (*n* = 7, age of 3 – 4 yrs, median age 3 yrs, male and female) was subcutaneously inoculated with 10^3^ PFU of RhCMV_UCD59_. Another group of animals (*n* = 12, age of 3 − 7 yrs, median age 3.5 yrs, all female) was used in a SIV vaccine study. Animals received either RhCMV_68-1_ or RhCMV_68-1_-derived RhCMV/SIV recombinant viruses, including RhCMV_68-1Gag_, RhCMV68_-1RTN_ and RhCMV_68-1Env_ (49) by a combination of subcutaneous inoculation (10^4^ PFU) and oral exposure (10^5^ PFU). Vaccination consisted of an initial priming at week 0 and two boosts at weeks 12 and 24. Starting on week 36, all animals received weekly low dose SIV challenges by the intravaginal route with SIV_mac251_ at a dose of 5 × 10^3^ TCID_50_ (0.54 ng/mL p27) until infected. Challenge SIV_mac251_ stock was kindly provided by N. Miller (NIH, Bethesda, MD) and Quality Biologics Inc. (Gaithersburg, MD). Blood samples collected from all animals were processed for plasma and peripheral blood mononuclear cells (PBMC) by Accu-Paque gradient centrifugation (Accurate Chemical & Scientific Corp., Westbury, NY) and cryopreserved at -80°C (plasma) or in liquid nitrogen (PBMC) for subsequent assessments.

### Serology Screening

Plasma IgG to a panel of specific pathogenic agents (SIV, SRV, STLV, BV, RhCMV, RRV, SFV) was measured on the Luminex (Austin, Tx) liquid bead based array platform using multiplex microbead bead reagents obtained from Charles River Laboratories (Wilmington, MA). By measuring the spectral properties of the beads and amount of associated R-phycoerythrin (R-PE), the median fluorescence intensity (MFI) for each specific antigen was determined. The sample MFI was compared against a previously validated known positive/negative cutoff value to qualitatively determine the presence or absence of antibody.

### Antibodies and Reagents

The following antibodies were purchased from the indicated manufacturers and used for flow cytometry; BD Bioscience: anti-CD3ε (SP34-2); Miltenyi Biotec: anti-NKG2A (REA110); MBL: anti-FcεR1γ (1D6); Beckman Coulter: anti-NKp46 (BAB281); Biolegend: anti-CD3ζ (6B10.2), anti-CD8 (RPA-T8), anti-CD16 (3G8), anti-GrB (GB11), anti-NKp30 (P30-15), anti-PECAM (WM59), anti-CD49f (GoH3), anti-CD2 (RPA-2.10), anti-SYK (4D10.2), anti-TNF-α (Mab11) and anti-IFN-γ (B27).

### Flow Cytometric Analysis of rhesus NK cells

Rhesus PBMCs and cell suspensions prepared from tissues were stained for flow cytometric analysis using fluorochrome-conjugated antibodies as previously described (11). Briefly, cell surface markers were stained with Abs and then fixed in 1.5% formaldehyde. To stain intracellular markers, samples were treated with permeabilization buffer containing 0.1% saponin, followed by staining of intracellular proteins. Analysis of PBMC samples collected from SPF animals with experimental viral infection was performed in a retrospective manner, while analysis of PBMC samples collected from non-SPF animals was performed in both retrospective and prospective manners.

### Functional analysis of rhesus NK cells

Telomerized rhesus fibroblast (Telo-RF) (50) were cultured in 96-well plates, infected with RhCMV_68-1_ (MOI = 1) for 2 hours, and then washed with PBS to remove unattached virus. At 3-4 days post infection, macaque PBMCs were added to the culture and incubated overnight in the presence of recombinant human IL-2 (10 U/ml). Six hours prior to analysis, 1µl autologous plasma was added along with Brefeldin A for cytokine analysis. For tumor and cytokine stimulation assays, macaque PBMCs were co-cultured for 12 hrs with human tumor target cells (K562 or 721.221) at a ratio of 10:1 (E:T) or 24 hrs with cytokines (human IL-12, IL-15 and IL-18). Six hours prior to analysis, Brefeldin A was added for analysis of cytokine expression. For CD16 crosslinking assays, macaque PBMCs were stimulated for 6 hrs with plate-bound anti-CD16 (3G8) antibody along with Brefeldin A (9). Following stimulation, NK cells were identified as CD3ε^−^ CD14^−^CD20^−/dim^NKG2A^+^ cell population.

### Statistics

Statistical analysis and graphing were conducted with Prism software (GraphPad Software, Inc., San Diego, CA). Wilcoxon matched-pairs signed-rank test was used for comparison of the frequency difference during NK cell functional assays. Nonparametric Mann-Whitney and chi-square tests were used for the difference in frequency and incidence of FcRγ^−^ NK cells between indicated groups, respectively. Correlation analyses were performed by Spearman’s rank tests. Differences were considered significant when p< 0.05.

## Acknowledgement

The authors would like to thank Abigail Spinner and Tracy Rourke (California National Primate Research Center, Davis, CA) for their help with flow cytometry; Drs.

Christopher J. Miller, Koen Van Rompay and Lauren A. Hirao (UC Davis) for providing macaque blood and tissue samples.

This work was supported in part by National Institutes of Health (NIH) grants AI110894 (to S.K.), DE011273 (to E.E.S. and P.A.B.), AI049342 (to P.A.B.), and ORIP/OD P51OD011132 (to Yerkes National Primate Research Center). Serology testing for this study was provided by the CNPRC Pathogen Detection Laboratory, which receives support from NIH grant P51OD011107.

The authors declare no competing financial interests.

## Author contributions

J.L. conceived the study, designed and performed experiments, interpreted data, and prepared the manuscript; W.C designed and performed experiments, conducted animal studies, interpreted data, and edited the manuscript; J.S. conceived the study, interpreted data, and prepared the manuscript; S.J., T.L., and P.P performed experiments; J. D. conducted animal studies; E. E. S., S. D., D.H., and P.B. directed the SPF-CMV infection work, interpreted data and edited the manuscript; S.K. conceived the study, interpreted data, supervised all aspects of the work, and prepared the manuscript.

