## Supplemental Figures for "FcRγ^−^ NK cell induction by specific CMV and expansion by subclinical viral infections in rhesus macaques"

### Supplemental Figure 1

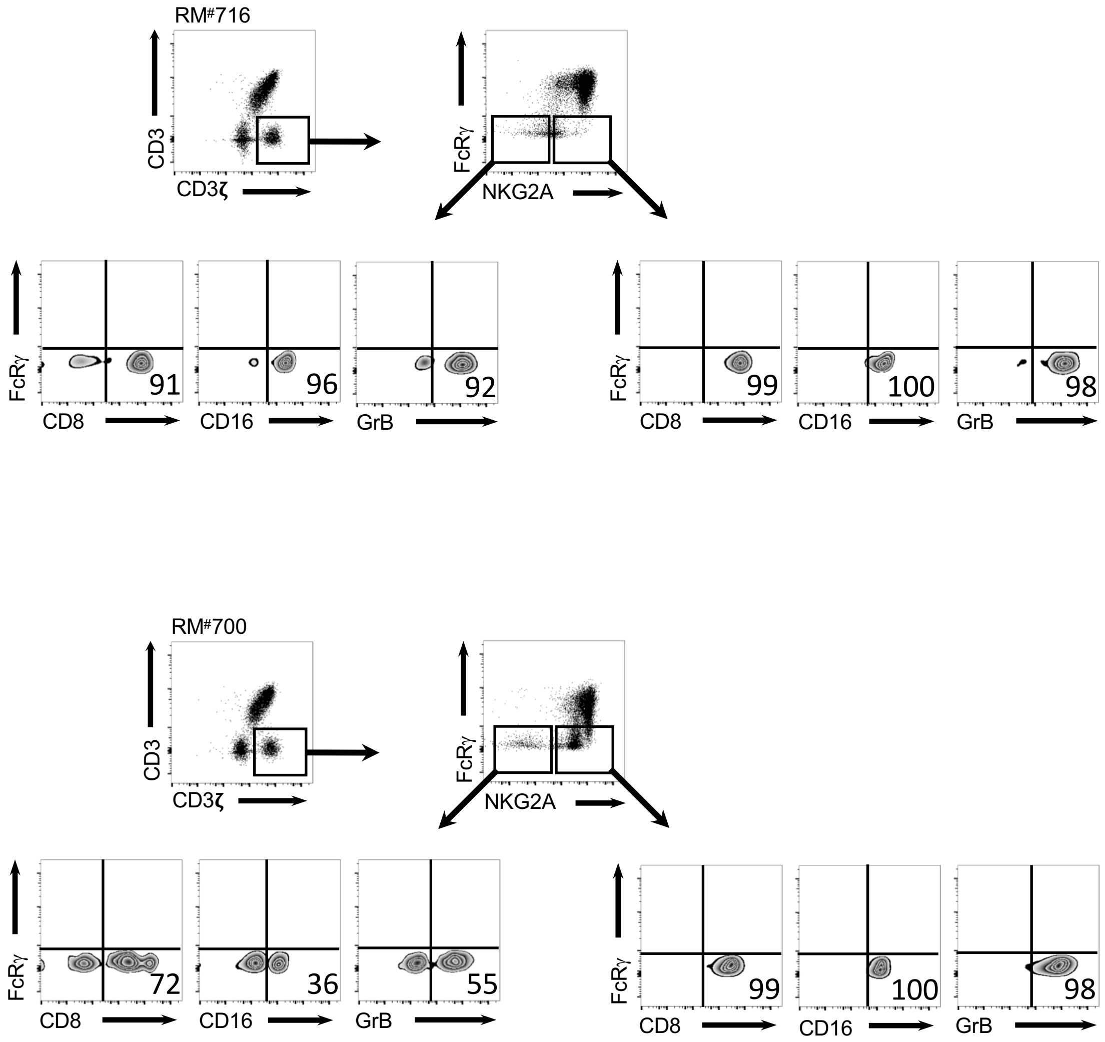


**Supplemental Figure 1. Comparative analysis of NKG2A$^+$ and NKG2A$^-$ FcRγ$^-$ subsets within the CD3ε$^-$CD14$^-$CD20$^{-/dim}$CD3ζ$^+$ population.** The CD3$^-$CD14$^-$CD20$^{-/dim}$CD3ζ$^+$FcRγ$^-$ cell population in PBMCs from two representative macaques was analyzed for the expression of NKG2A, and expression of indicated NK cell-associated markers was compared between the NKG2A$^+$ and NKG2A$^-$ subsets.

### Supplemental Figure 2

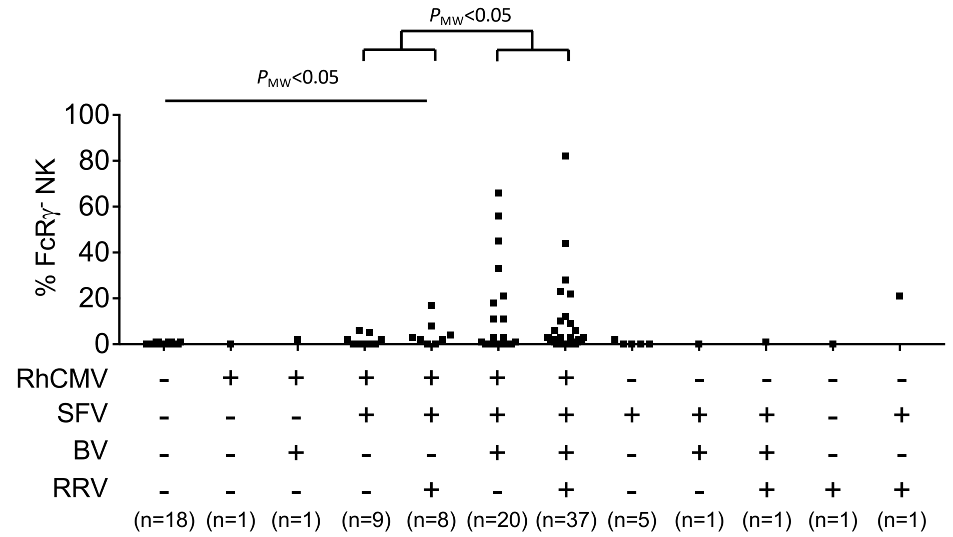


**Supplemental Figure 2. Association of FcRγ$^-$ NK cells with infection by RhCMV, SFV, BV and RRV.** Frequencies of FcRγ$^-$ NK cells within individual macaques grouped according to their IgG serological status with respect to 4 different common viruses (RhCMV, SFV, BV and RRV). Statistical analyses were performed using nonparametric Mann-Whitney tests.
